## Supplementary Figures for "Aberrant cortical activity, functional connectivity, and neural assembly architecture after photothrombotic stroke in mice"

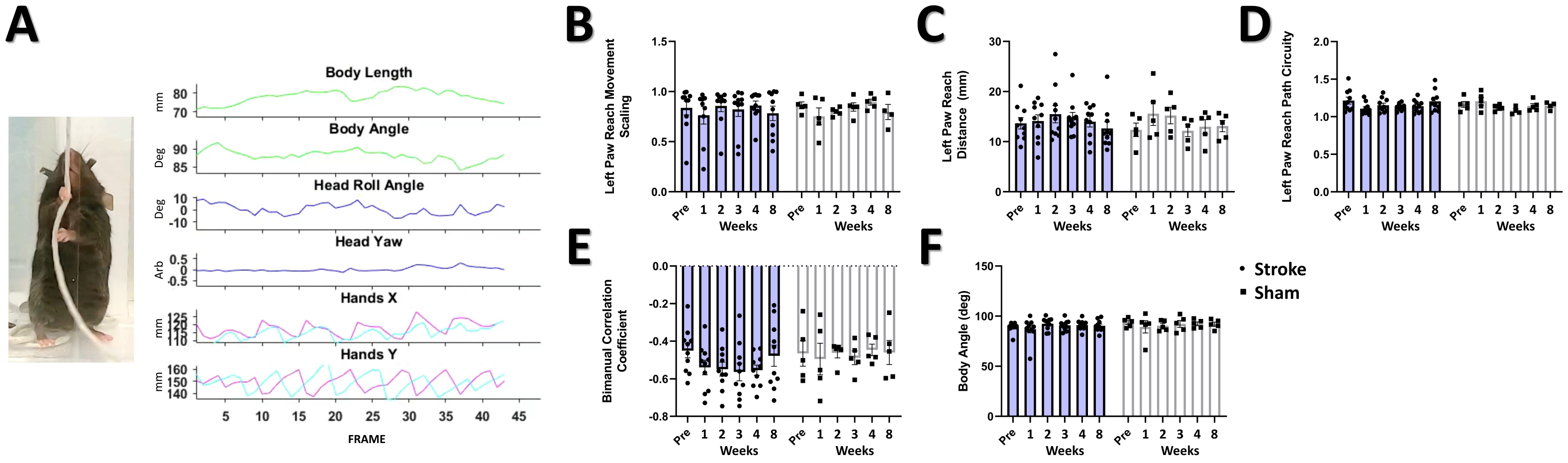
A
Body Length
Body Angle
Head Roll Angle
Head Yaw
Hands X
Hands Y
mm
Deg
Arb
FRAME
B
Left Paw Reach Movement Scaling
C
Left Paw Reach Distance (mm)
D
Left Paw Reach Path Circuity
E
Bimanual Correlation Coefficient
F
Body Angle (deg)
Pre
Weeks
Stroke
Sham

**Fig S1. String pull task does not detect a behavioral deficit after stroke.** (**A**) Left: Image of example animal performing the string pull task at the pre-stroke time. Right: Data for the animal's body length, body angle, head angle and yaw, and position of the hands is tracked to determine alterations in motor movement across times. No significant main effect or interaction are seen in the left (affected) paw reach movement scaling (**B**), reach distance, (**C**), reach path circuity (**D**), bimanual correlation coefficient (**E**), or in the animal body angle (**F**). Stroke N = 10, Sham N = 5. $^*p < 0.05$; $^{**}p < 0.01$; $^{***}p < 0.001$

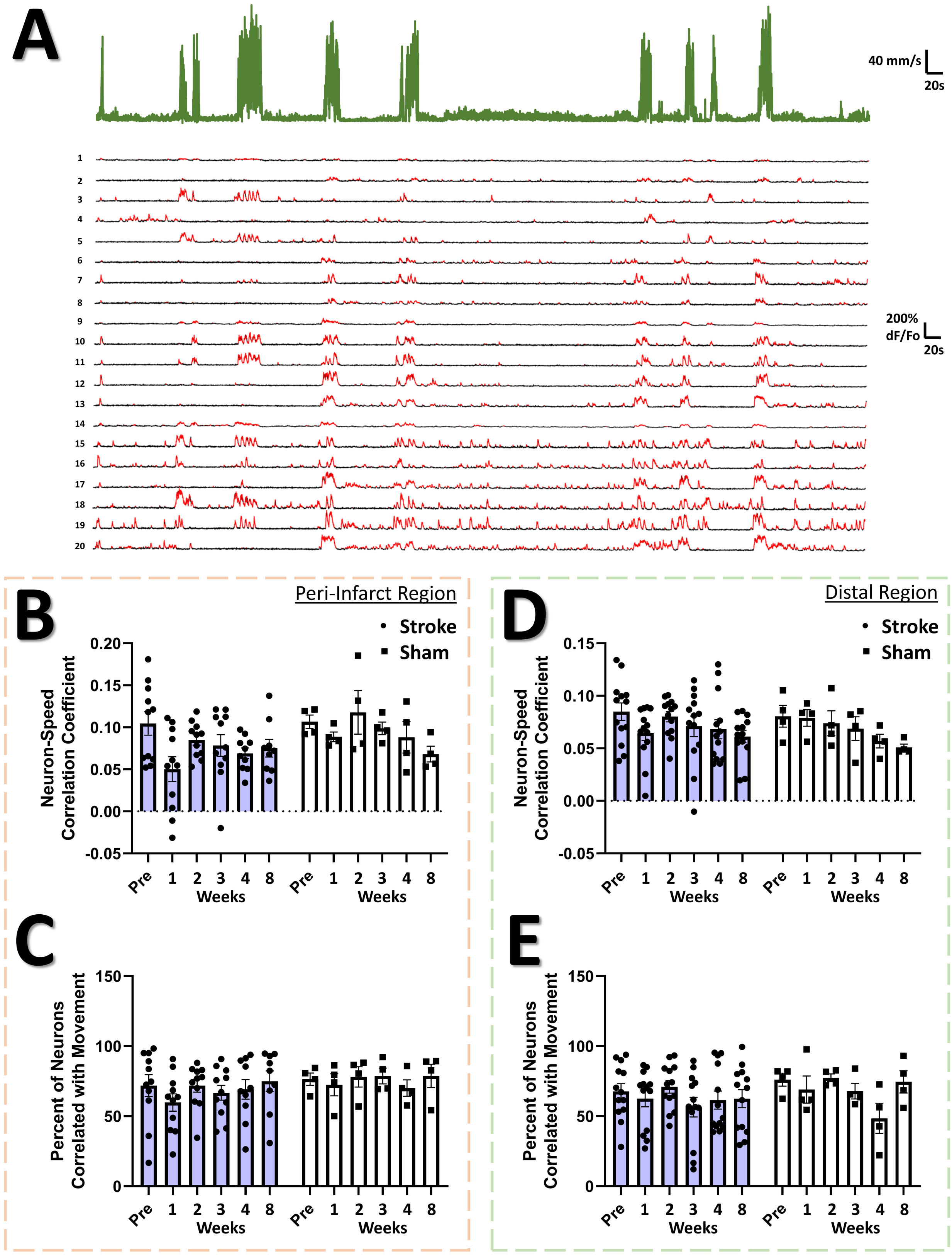
A
40 mm/s
20s
200% dF/Fo
20s
B
Peri-Infarct Region
Stroke
Sham
Neuron-Speed Correlation Coefficient
Weeks
C
Percent of Neurons Correlated with Movement
D
Distal Region
E
Pre
Weeks

55

**Fig S2. Average neuron correlation with animal movement is not detectably altered by stroke.** (**A**) Example time series of animal speed (green) and $Ca^{2+}$ traces from the 20 most active neurons of the population co-recorded with the animal movement speed. Significant $Ca^{2+}$ transients are highlighted by red segments. For the peri-infarct imaging region, a significant main effect of time was observed in the average correlation coefficient between neuron $Ca^{2+}$ traces and speed (**B**), however *post-hoc* tests did not show a significant difference between any of the times within the stroke and sham groups, respectively. Mixed Effects Model, Time $F_{(3.116, 38.64)} = 2.861$, $P = 0.0474$; Stroke Group $F_{(1, 13)} = 2.296$, $P = 0.1536$; Interaction $F_{(5, 62)} = 0.7734$, $P = 0.5726$. No main effects or interaction were found in the percent of neurons correlated with movement for the peri-infarct imaging region (**C**). In the distal imaging region, no significant main effects or interactions were seen in the average correlation coefficient between neuron $Ca^{2+}$ traces and speed (**D**) or in the percent of neurons correlated with movement (**E**). Peri-infarct region Stroke N = 11, Sham N = 4. Distal region Stroke N = 13, Sham N = 4. *$p < 0.05$; **$p < 0.01$; ***$p < 0.001$

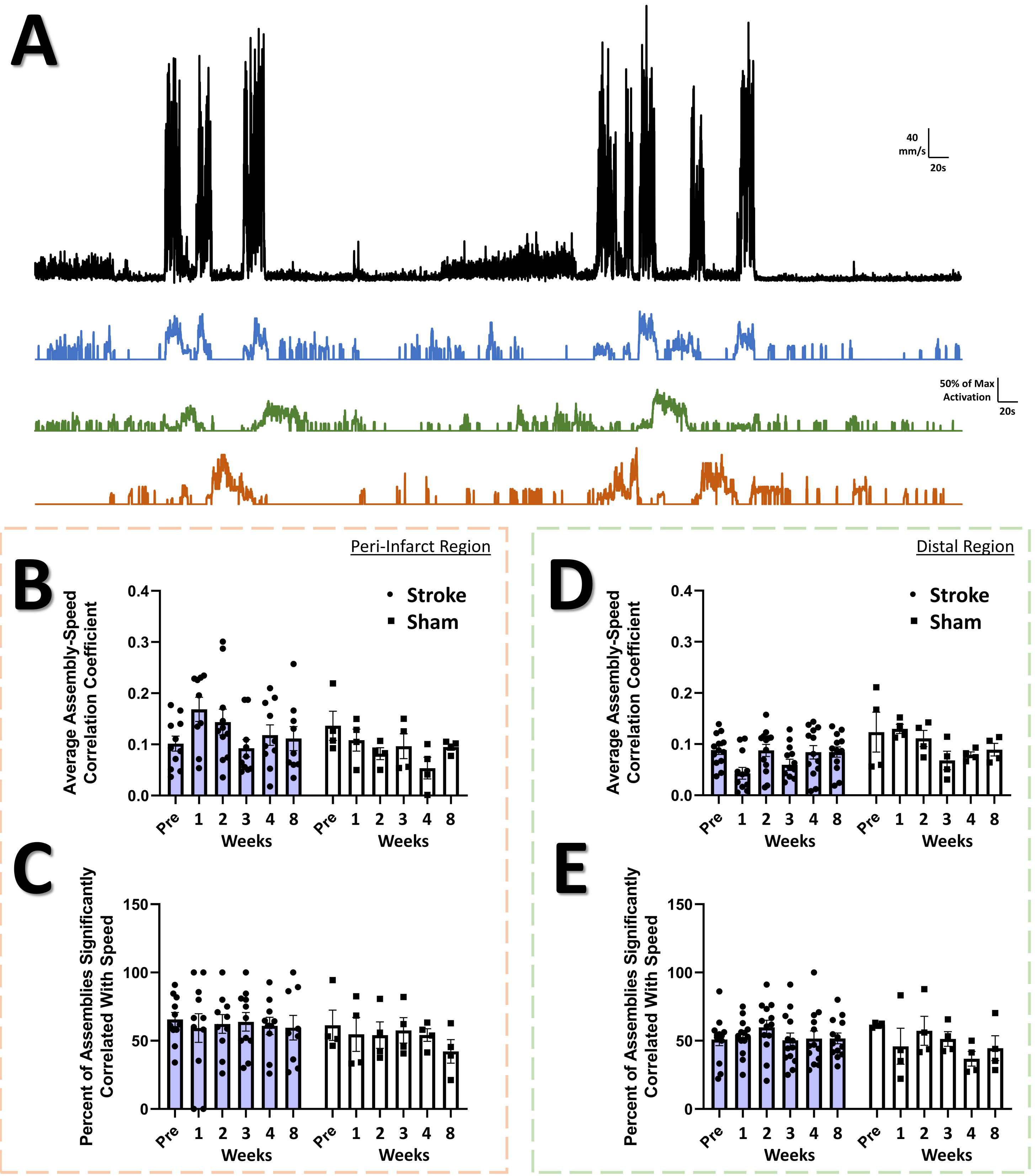
A
40
mm/s
20s
50% of Max
Activation
20s
Peri-Infarct Region
B
Stroke
Sham
Average Assembly-Speed
Correlation Coefficient
0.0
0.1
0.2
0.3
0.4
Pre 1 2 3 4 8
Weeks
Pre 1 2 3 4 8
Weeks
C
Percent of Assemblies Significantly
Correlated With Speed
0
50
100
150
Pre 1 2 3 4 8
Weeks
Pre 1 2 3 4 8
Weeks
Distal Region
D
Stroke
Sham
Average Assembly-Speed
Correlation Coefficient
0.0
0.1
0.2
0.3
0.4
Pre 1 2 3 4 8
Weeks
Pre 1 2 3 4 8
Weeks
E
Percent of Assemblies Significantly
Correlated With Speed
0
50
100
150
Pre 1 2 3 4 8
Weeks
Pre 1 2 3 4 8
Weeks

**Fig S3. Correlation between assembly activations and movement is not detectably affected by stroke.** (**A**) Representative time series of animal movement speed (black) and the assembly activation level for three representative assemblies (blue, green, orange) from the neuron population corresponding to the animal movement speed above. In the peri-infarct imaging region, no main effects or interactions are seen in the average assembly-speed correlation coefficient (**B**), or in the percent of assemblies significantly correlated with speed (**C**). In the distal imaging region, no main effects or interactions are seen in the average assembly-speed correlation coefficient (**D**), or in the percent of assemblies significantly correlated with speed (**E**). Peri-infarct region Stroke $N = 11$, Sham $N = 4$. Distal region Stroke $N = 13$, Sham $N = 4$. *$p < 0.05$; **$p < 0.01$; ***$p < 0.001$
